# Development of a New Method for the Highly Effective Identification of Cold Resistance in Living Avocado Varieties

**DOI:** 10.1101/481671

**Authors:** Zheng Chunlin, Wu Fan, Li Cuiling, Peng Wenli, Zhang Shaofeng, Zhu Jingjie, Jiang Hao, Li Maofu

## Abstract

This paper first identified the cold resistance of 38 varieties of avocado by determining the semi-lethal low temperature (LT_50_) of the leaves using an electrical conductivity method in combination with a logistic function, and then analyzed the correlation between the LT_50_ of 27 varieties and the capacitance measured 9 different parts of the leaves in vivo, to explore the relationship between the cold resistance of various avocado varieties and the capacitance of different parts of avocado leaves, so as to develop a new method for highly effective identification of cold resistance of living avocado varieties. The results showed that various avocado varieties’ LT_50_ was significantly positively correlated with the capacitance of some parts of leaves, showing that the cold resistance of various avocado varieties was negatively correlated with the capacitance of various leaf parts. The results of mango variety trials conducted for comparison is coincident with the theoretical conclusion reached in the identification of the cold resistance of the avocado. So as to the study of the cold resistance of avocado and mango varieties, the capacitance of live mature leaves measured in the field can be used as a new method for the judgment of cold resistance.

**Highlight:** A new method for identification of avocado cold resistance through measuring the capacitance of different parts of leaves was developed. This method are simple, quick and efficient.

## Introduction

The avocado (*Persea americana Mill*) is a fast-growing evergreen arbor fruit tree of the genus Persea in the laurel family. Native to the tropical humid areas in Central America and Mexico as well as high-attitude mountainous forests or tropical plateaus, the avocado is a well-known tropical and subtropical fruit tree (***Ge Yu et al., 2017***). The avocado was introduced into China in 1918, and now is planted in tropical regions including Hainan, Guangdong, Guangxi, Guizhou, Yunan and Taiwan. Low temperature is a key limiting factor for fruit farming (***Dong Meichao et al., 2016***), so the avocado suffers severe chilling damage in winter in subtropical high-elevation regions such as North Guangxi and Guizhou (**Lv Shihong et al., 1997**). Its leaves may suffer injury, and in severe cases, the whole plant may die. This seriously affects the secure development of China’s avocado industry. Therefore, it is of great theoretical and practical significance to identify the cold resistance and tolerable low temperature range of avocado varieties, in order to guide regional avocado farming, cold resistant variety breeding and in-depth explorations of the cold resistance mechanism.

The electrical conductivity method is a classical method for the identification of the cold resistance of plant tissues, and widely used due to its ease and high efficiency (***Bu Qingyan et al., 2005***). On that basis, the “guide line” of plant tissues at varying temperature can be determined, and the inflection temperature can be measured using a logistic function to calculate the LT_50_ of plant tissues as a quantitative index related to plant cold resistance (***Zhu Genhai et al., 1986***). In recent years, the electrical conductivity method has been widely applied to the research and identification of the cold resistance of fruit trees such as apple (***Yang Fengqiu et al., 2011; Shi Chao et al., 2013***), grape (***Lu Jinxing et al.,2012; Zhang Qian et al., 2013***), pear (***Li Juncai et al., 2007***), syzygium samarangense (***Zhang Lvpeng et al., 2012***), *Prunus mume* plum (***Gao Hongzhi et al., 2005***), *Prunus salicina* plum (**Liu Weisheng et al., 1999**), orange (***Luo Zhengrong et al., 1992***) and carambola (***Ren Hui et al., 2016***), but there is no report on the combined use of the electrical conductivity method with a logistic equation to determine the LT_50_ and cold resistance of the avocado.

Due to the accuracy and practicability of the electrical conductivity method or LT_50_ identification method, its use requires a simulation of low temperature stress and cutting of plant tissues. To avoid this destructive identification method, it is necessary to develop a method for identification in vivo to identify the cold resistance of avocado plants accurately and efficiently.

Capacitance is a physical quantity used to represent the charge capacity of a capacitor. Its value, related to the dielectric constant for the material, the measuring area and the inter-pole distance (***D. Halliday et al., 2002***), can be directly measured with a capacitance meter. Given a measuring area and inter-pole distance, the value of the capacitance relates only to the dielectric constant of the observed samples. Given samples, the capacitance measured with a given capacitance meter should be a fixed value, and this makes it possible to infer sample difference based on the capacitance measured with the given meter. However, according to our data, there is currently no study on the inference of plant sample difference by capacitance mensuration.

In this paper, therefore, we combined the electrical conductivity method with a logistic function to measure the LT_50_ of 38 avocado varieties from Avocado Varieties Nursery, College of Tropical Agriculture and Forestry, Hainan University. Meanwhile, we measured the capacitance in different parts of 27 varieties of in vivo avocado leaves, and made an analysis of the correlation between the LT_50_ and capacitance of each part in order to determine whether the electrical conductivity method could be used to identify the cold resistance of avocado varieties. Then, we measured the capacitance of the leaves of two big mango varieties: Tainong No. 1 and Dongmang in the field in order to further validate the reliability of the electrical conductivity method for the identification of plant cold resistance.

## 1 Materials and Methods

### 1.1 Test Materials

The test materials came from Avocado Varieties Nursery, College of Tropical Agriculture and Forestry, Hainan University, including 38 avocado varieties, each from three healthy sexennially bearing trees. Samples were collected in December 2017 and January 2018: 18 healthy, intact mature leaves of basically the same size were collected from the four cardinal sides (i.e. east, south, west and north) of every tree. Capacitance for nine of the 18 was measured in vivo, while the remaining nine were wrapped in wet gauze and taken back to the laboratory for relative electrical conductivity testing.

### 1.2 Test Methods

#### 1.2.1 Disposal of Low-temperature Stress

The 38 portions of avocado leaves were cleaned with running water and then washed twice with distilled water. After the surface was wiped dry with filter paper, the leaves were dark-treated for 12 hours at 4^°^C, 2^°^C, 0^°^C, −2^°^C, −4^°^C, −6^°^Cand −8^°^C respectively, with treatment at room temperature (25^°^C) as control. In this paper, the varieties were numerically numbered. See Appendix 1 for detailed information on the varieties.

#### 1.2.2 Relative electrical conductivity measurement

After treatment, we took out the material and put it in a preservation box, and put the preservation box in an ice water mixture of 0^°^C (an ice water mixture was not used on the room temperature, 4^°^C, 2^°^C or 0^°^C samples) to unfreeze it for 30min. Then we cut the leaves into pieces of about 1cm^2^, avoiding larger veins. The samples for each variety of leaves treated at each individual temperature were combined into 304 groups (eight temperature treatments for each of 38 varieties). From each group, we took six 0.5g samples and put them in ten 100ml Erlenmeyer flasks (the flasks were cleaned beforehand with dishwashing liquid, washed with distilled water and finally dried). Then we put 25ml of distilled water in each Erlenmeyer flask and placed it in a vacuum drying oven to exhaust air for 30min, and placed it on a shaking table to shake for 1h. The flasks were then removed from the oven and left still for 15min, after which electrical conductivity K1 was measured with a portable electrical conductivity meter (Manufacturer: Hangzhou Qiwei Instrument Co., Ltd, model: DDB-11A), then each flask was weighed. The flasks were then immersed in a boiling bath and removed for 15min of cooling, after which water was added to restore the flask weight. The flasks were shaken on the shaking table for one hour, removed and allowed to remain still for 10min, and finally K2 was measured.

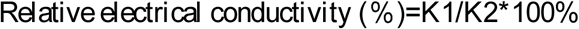

#### 1.2.3 Logistic Equation Fitting and Inflection Temperature Calculation

By reference to ***Zhu Genhai et al.(1986)***’s method, we used the logistic function to determine the LT_50_ of plant tissues. In other words, we used SPSS19 to make a regression analysis of the electrolyte leakage curve of each variety treated at various low temperatures by fitting the logistic function (y=k/(1+a*exp(−bx)). The calculation method used for inflection temperature was LT_50_=(*ln*a)/b.

#### 1.2.4 Capacitance Mensuration

We used a capacitance meter (Chinese brand Mastech, model MS8910) to test the capacitance in the main vein at the leaf stalk base, the main vein at the middle leaf section, the main vein at the leaf tip, the lateral vein at the leaf stalk base, the lateral vein at the middle leaf section, the lateral vein at the leaf tip, the mesophyll in the leaf stalk base, the mesophyll in the middle leaf section and the mesophyll in the leaf tip (see Fig.1). During testing, we inserted the capacitance meter’s metal testing plate into the site to be measured to assess the capacitance.

**Fig 1:**
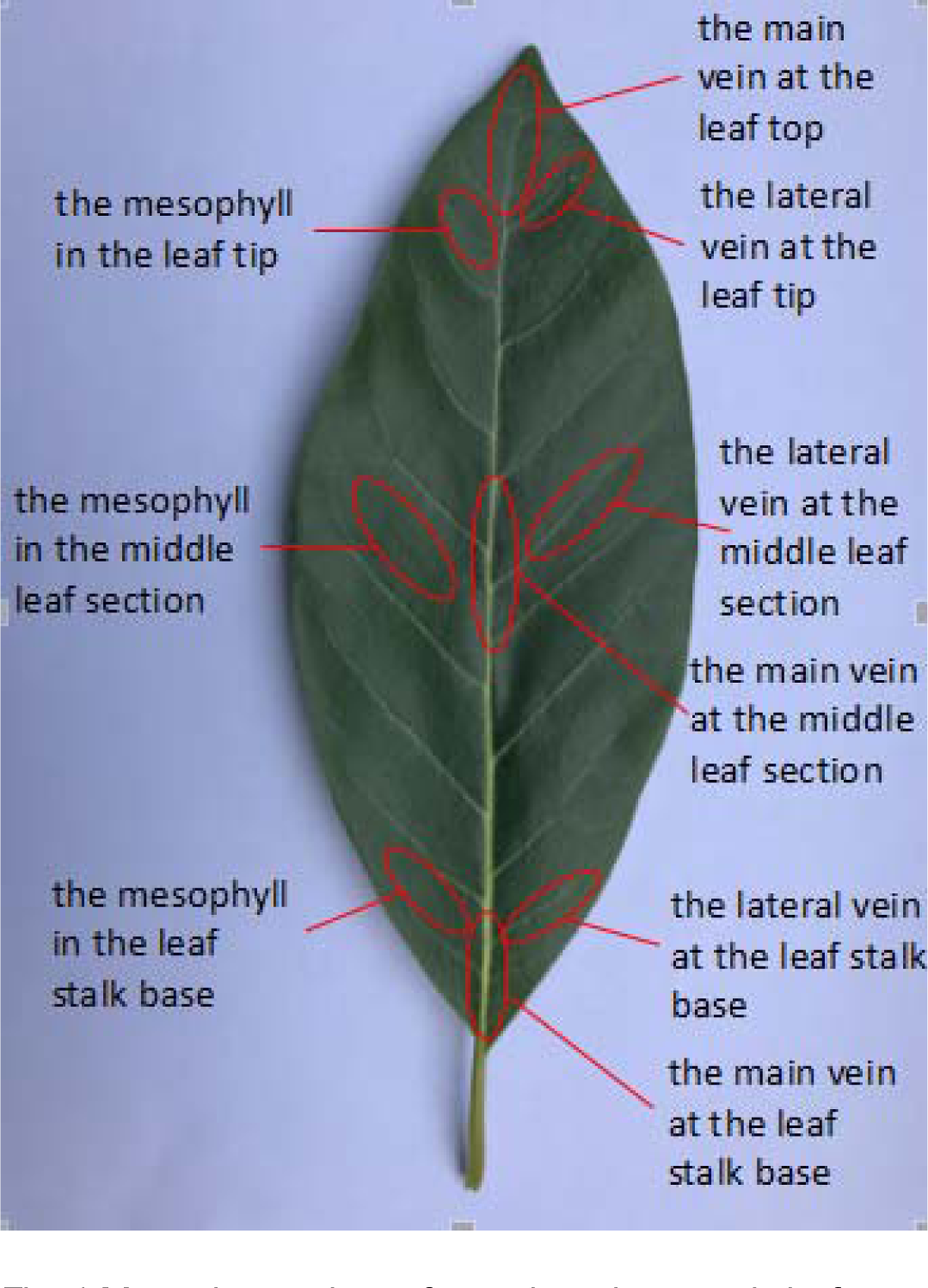
Measuring sections of capacitanc in avocado leaf.

### 1.3 Confirmatory trials on mango cold resistance

Dongmang (*Mangifera hiemalis Liang Jian-Ying*) and Tainong No. 1 (*Mangifera indica cv Tainong*), two mango varieties with significantly different cold resistance, were selected from Mango Varieties Nursery, Genetic Resources Institute, Chinese Academy of Tropical Agricultural Sciences, for confirmatory trials on the cold resistance of capacitance (***Wu Zehuan et al., 1984; Huang yun et al., 2013***). Dongmang showed higher cold resistance than Tainong No. 1. After selecting ten healthy, intact and mature leaves sprouted in the current year of either variety from three octennially bearing trees, we measured the capacitance in the main vein at the leaf stalk base, the main vein at the middle leaf section, the main vein at the leaf tip, the lateral vein at the leaf stalk base, the lateral vein at the middle leaf section, the lateral vein at the leaf tip, the mesophyll in the leaf stalk base, the mesophyll in the middle leaf section and the mesophyll in the leaf tip.

## 2 Results and Analyses

### 2.1 Analysis of the relative electrical conductivity of different avocado leaves under low-temperature stress

As can be seen in Table 1, the relative electrical conductivity of avocado leaves is at the minimum level at ambient temperature, but there was a significant difference among the 38 avocado varieties. Variety No. 30 showed electrical conductivity of only 12.58%, significantly lower than the other varieties, while No. 27 had the highest electrical conductivity, up to 12.56% higher than that of No. 30. After low-temperature treatment, the relative electrical conductivity of the avocado varieties increased to varying degrees, but there was a difference in the amplification of relative electrical conductivity among the avocado varieties at the same low temperature. Specifically, after treatment at 4 ^°^C ∼-6 ^°^C, the relative electrical conductivity of No. 1 and No. 14 avocado varieties increased by 37.35% and 40.05% respectively, while the relative electrical conductivity of No. 28 increased by 68.56% after treatment at 4^°^C∼-6^°^C, and that of No. 43 increased by 68.62% after treatment at 2^°^C∼-8^°^C. This shows a significant difference in relative electrical conductivity among the 38 avocado varieties at room temperature, and also a big difference in the range of relative electrical conductivity under the same stress conditions. In other words, different avocado leaves have different sensitivities to low temperature, but there is no absolute relation between the electrical conductivity of a leaf and its sensitivity to low temperature.

**Table 1:**
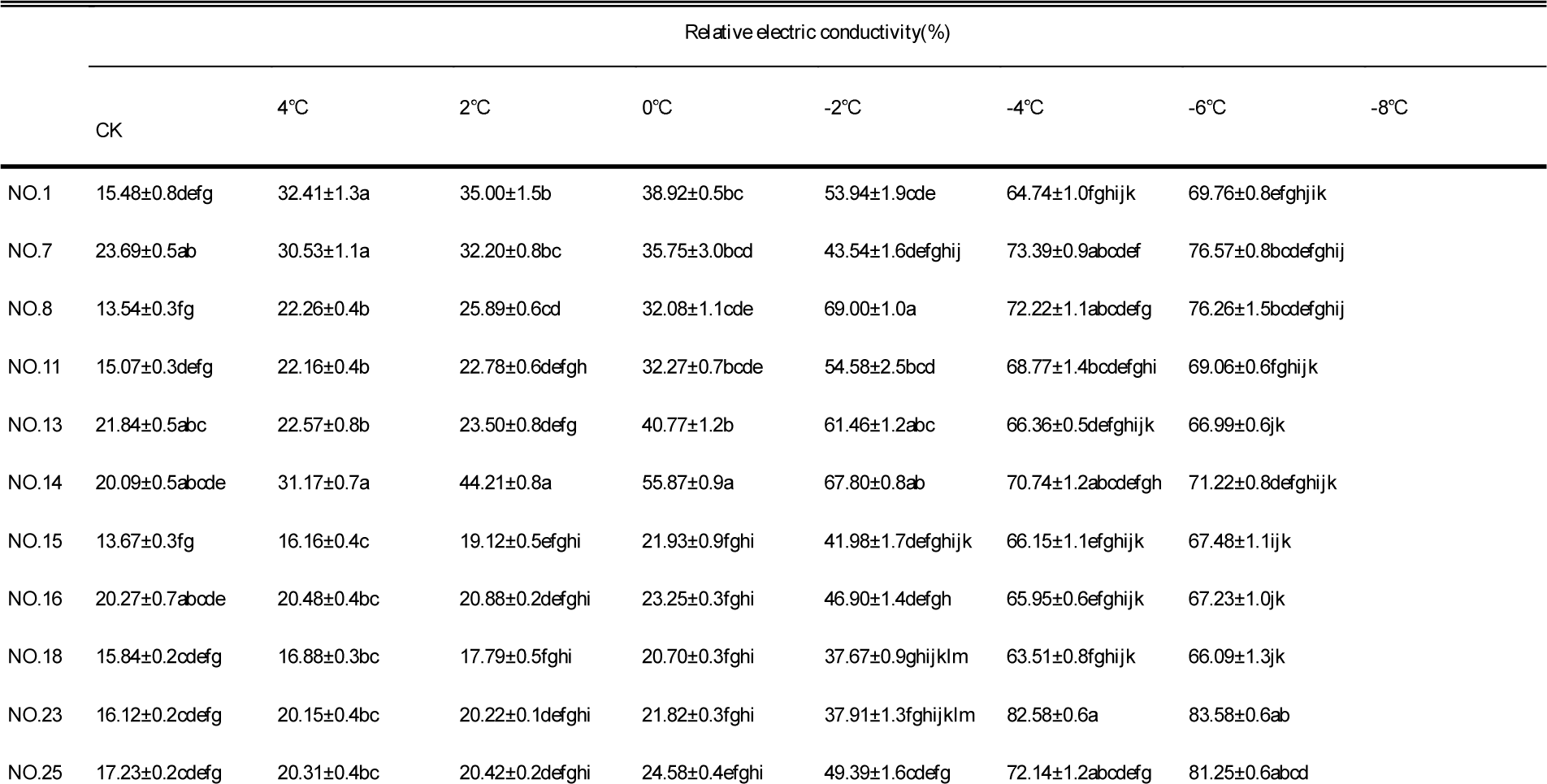
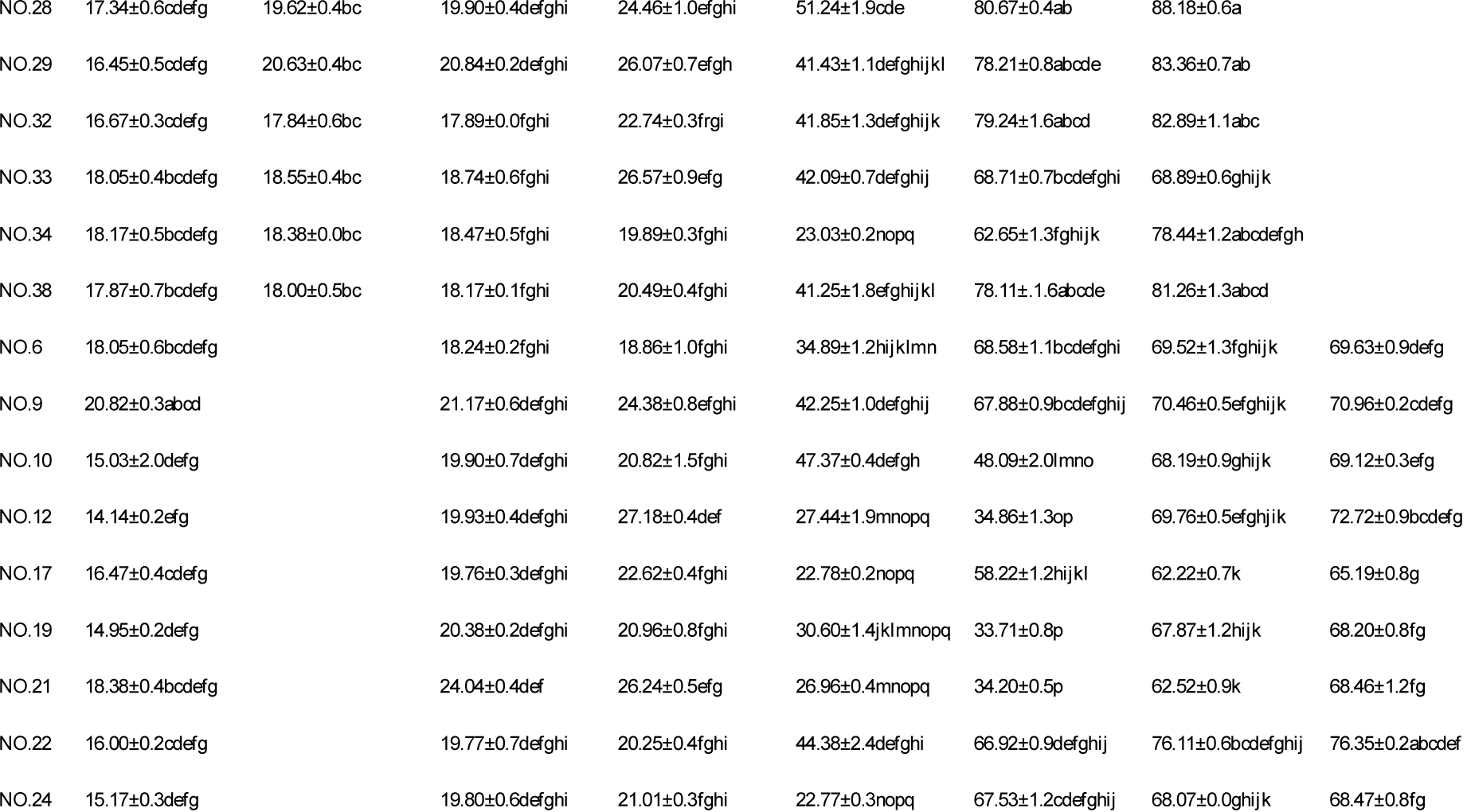
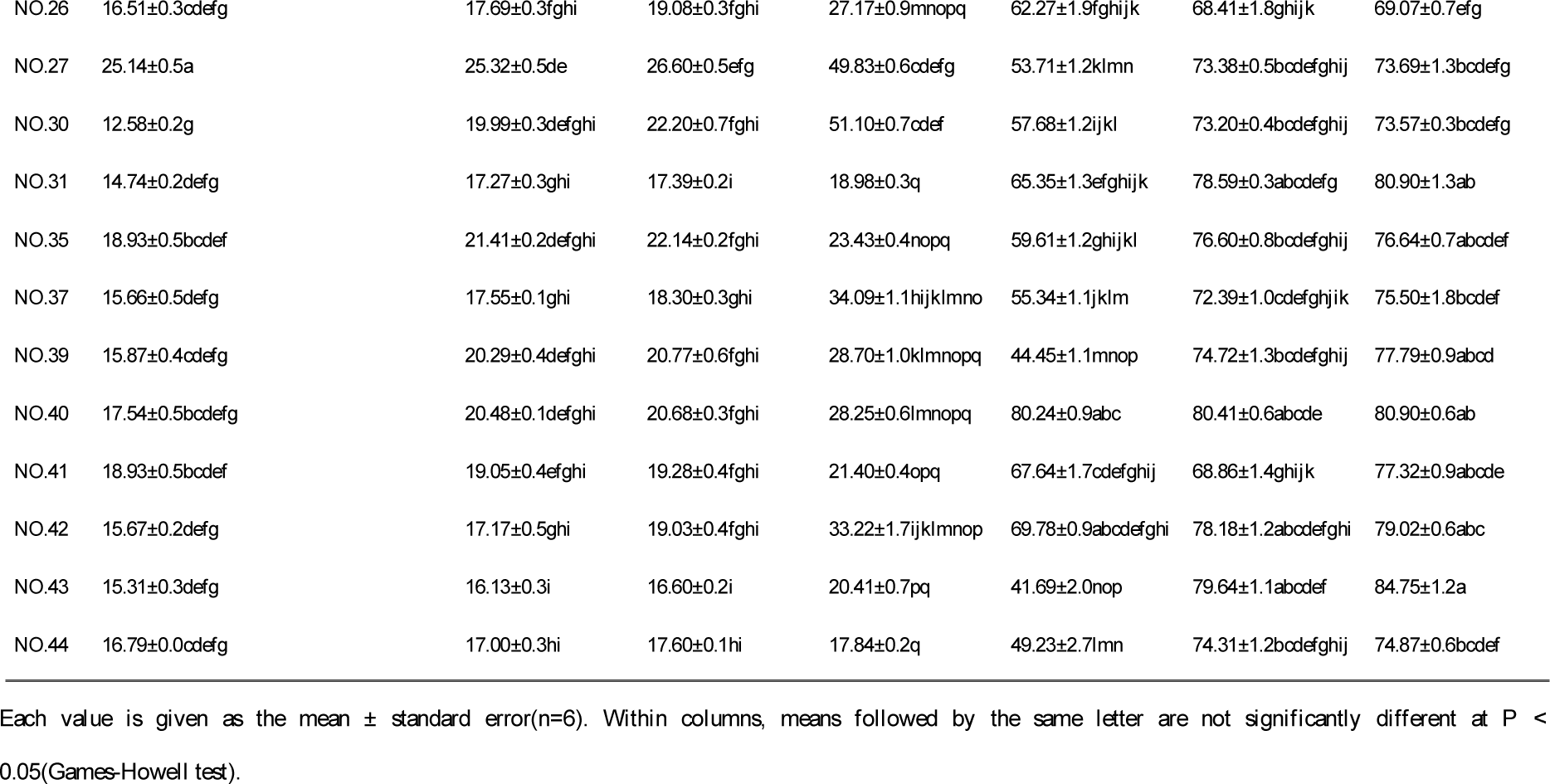
Relative electrical conductivity of 38 avocado varieties leaves under different temperatures stress.

### 2.2 Analysis of LT_50_ for different avocado varieties under low-temperature stress

For ease of data analysis, we divided the relative electrical conductivity curves of the 38 avocado varieties into two groups according to the low temperature range and performed a diagraph analysis (Fig.2a, 2b). As can be seen in Fig.2, the relative electrical conductivity of the 38 avocado varieties showed a relative rising trend with the temperature falling, and this conforms to the change characteristics of the “guide line”. The inflection temperature of 17 varieties appeared between 2^°^C and −4^°^C (Fig.2a), while that of 21 varieties appeared between 0^°^C and −6^°^C (Fig.2b).

**Fig 2:**
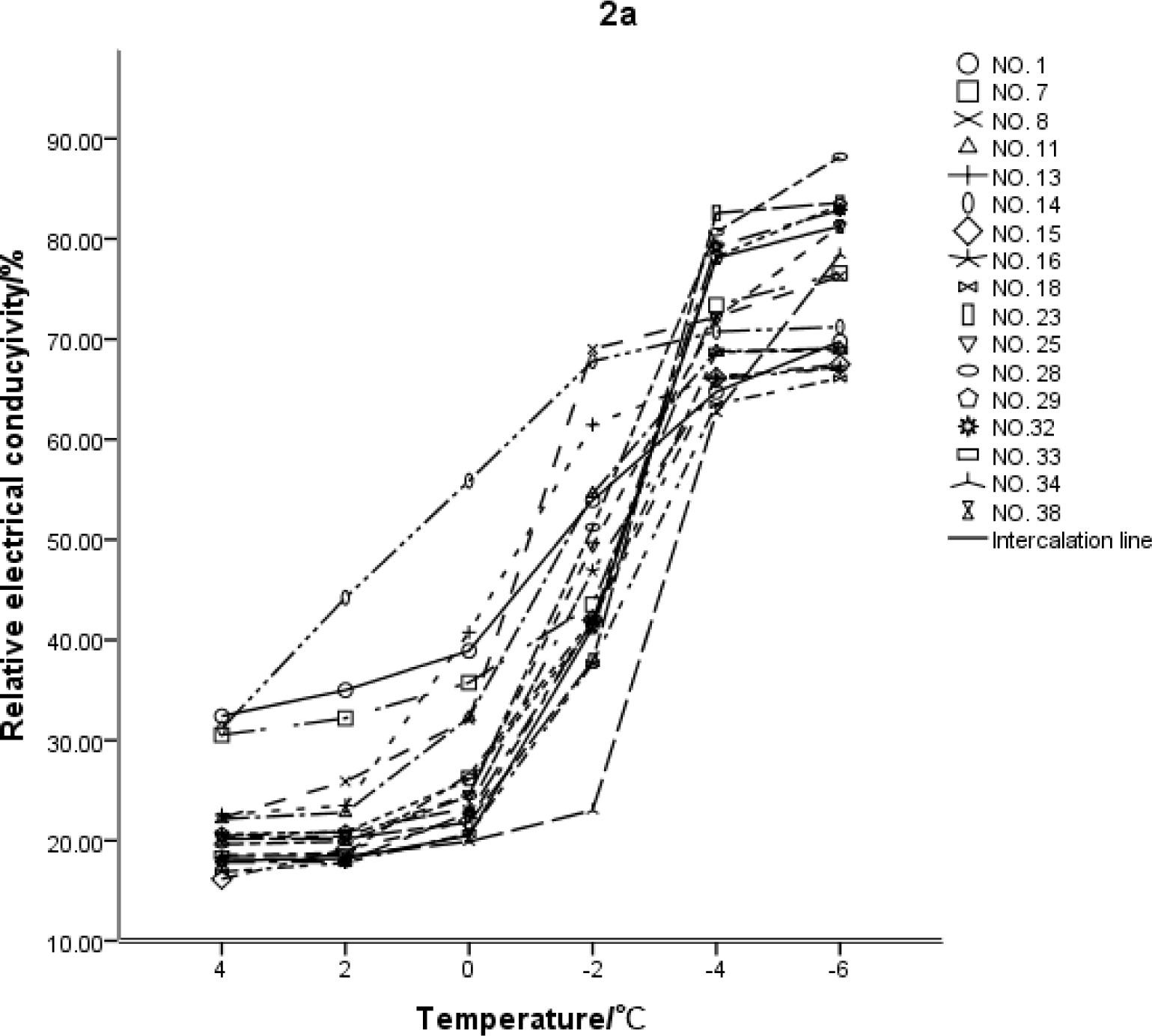
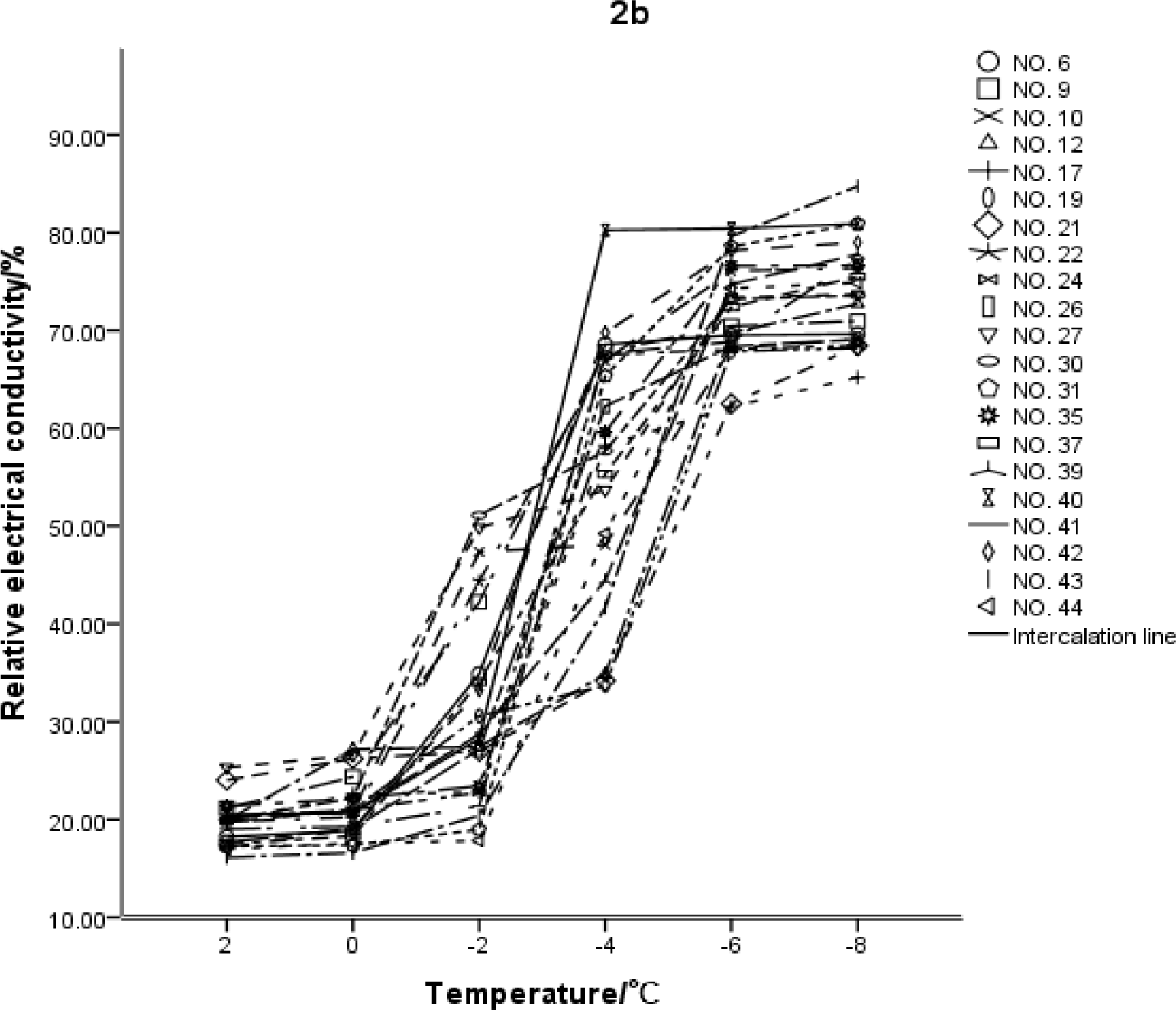
Varying curves of relative electrical conductivity of avocado varieties at different temperatures.

Table 2 shows the results of a regression analysis on the relative electrical conductivity curve of the 38 avocado varieties conducted by fitting the logistic function. Number 14 had the highest LT_50_ at 3.045^°^C, and No. 21 had the lowest LT_50_ at −5.056^°^C, while the others’ ranged from −5.056^°^C to 0.854^°^C. The value of LT_50_ can help identify the cold resistance of the various varieties: the lower the LT_50_ value, the higher the cold resistance. This shows that of the 38 avocado varieties, No. 21 had the strongest cold resistance, while No. 14 had the weakest cold resistance.

**Table 2:** Logistic equation and semi-lethal low temperature of 38 varieties of avocado.

|  | Equation | R<sup>2</sup> | LT<sub>50</sub>(°C) |
| --- | --- | --- | --- |
| No. 1 | $y=100/(1+1.247\exp(0.177x))$ | 0.955 | -1.247 |
| No. 6 | $y=75.644/(1+2.408\exp(0.545x))$ | 0.927 | -1.612 |
| No. 7 | $y=100/(1+1.375\exp(0.229x))$ | 0.868 | -1.391 |
| No. 8 | $y=88.019/(1+1.045\exp(0.376x))$ | 0.913 | -0.117 |
| No. 9 | $y=77.302/(1+1.671\exp(0.458x))$ | 0.951 | -1.121 |
| No. 10 | $y=78.991/(1+1.884\exp(0.347x))$ | 0.936 | -1.825 |
| No. 11 | $y=86.893/(1+1.269\exp(0.312x))$ | 0.946 | -0.764 |
| No. 12 | $y=100/(1+3.514\exp(0.276x))$ | 0.881 | -4.553 |
| No. 13 | $y=74.496/(1+0.713\exp(0.396x))$ | 0.950 | 0.854 |
| No. 14 | $y=74.665/(1+0.304\exp(0.391x))$ | 0.993 | 3.045 |
| No. 15 | $y=94.526/(1+2.238\exp(0.32x))$ | 0.941 | -2.517 |
| No. 16 | $y=98.157/(1+2.031\exp(0.278x))$ | 0.921 | -2.549 |
| No. 17 | $y=78.868/(1+2.474\exp(0.348x))$ | 0.877 | -2.603 |
| No. 18 | $y=100/(1+2.654\exp(0.298x))$ | 0.933 | -3.275 |
| No. 19 | $y=100/(1+3.646\exp(0.266x))$ | 0.891 | -4.863 |
| No. 21 | $y=100/(1+3.215\exp(0.231x))$ | 0.872 | -5.056 |
| No. 22 | $y=82.64/(1+2.155\exp(0.491x))$ | 0.966 | -1.564 |
| No. 23 | $y=138.358/(1+3.631\exp(0.313x))$ | 0.894 | -4.120 |
| No. 24 | $y=77.887/(1+2.72\exp(0.465x))$ | 0.847 | -2.152 |
| No. 25 | $y=130.189/(1+2.948\exp(0.28x))$ | 0.955 | -3.861 |
| No. 26 | $y=77.592/(1+2.841\exp(0.471x))$ | 0.922 | -2.217 |
| No. 27 | $y=88.095/(1+1.698\exp(0.296x))$ | 0.948 | -1.789 |
| No. 28 | $y=125.323/(1+2.764\exp(0.337x))$ | 0.953 | -3.017 |
| No. 29 | $y=99.783/(1+2.017\exp(0.353x))$ | 0.926 | -1.988 |
| No. 30 | $y=79.152/(1+1.719\exp(0.435x))$ | 0.959 | -1.245 |
| No. 31 | $y=89.855/(1+5\exp(0.545x))$ | 0.913 | -2.953 |
| No. 32 | $y=119.922/(1+3.083\exp(0.352x))$ | 0.935 | -3.199 |
| No. 33 | $y=99.438/(1+2.173\exp(0.297x))$ | 0.939 | -2.613 |
| No. 34 | $y=100/(1+3.609\exp(0.391x))$ | 0.860 | -3.282 |
| No. 35 | $y=100/(1+2.794\exp(0.299x))$ | 0.869 | -3.436 |
| No. 37 | $y=88.196/(1+3.059\exp(0.396x))$ | 0.976 | -2.823 |
| No. 38 | $y=117.95/(1+3.13\exp(0.354x))$ | 0.926 | -3.223 |
| No. 39 | $y=100/(1+3.804\exp(0.334x))$ | 0.940 | -4.000 |
| No. 40 | $y=96.207/(1+2.585\exp(0.404x))$ | 0.852 | -2.351 |
| No. 41 | $y=86.414/(1+3.561\exp(0.46x))$ | 0.882 | -2.761 |
| No. 42 | $y=97.965/(1+2.663\exp(0.358x))$ | 0.921 | -2.736 |
| No. 43 | $y=105.714/(1+5\exp(0.376x))$ | 0.930 | -4.280 |
| No. 44 | $y=92.549/(1+5\exp(0.423x))$ | 0.921 | -3.805 |

### 2.3 Correlation between LT_50_ and the electrical conductivity of avocado varieties at different temperatures

According to Pearson’s correlation analysis of the LT_50_ of the avocado varieties and their relative electrical conductivities at different temperatures (Table 3), there is a highly significant positive correlation between LT_50_ and the relative electrical conductivity at 4^°^C, 2^°^C, 0^°^C, −2^°^C and −4^°^C, while there is an insignificant correlation between LT_50_ and the relative electrical conductivity at −6^°^C and −8^°^C and control temperature. As can be seen in Table 1, at −4^°^C, the relative electrical conductivity of 82% of the avocado varieties tested was greater than 50%; at −6^°^C, the relative electrical conductivity of all the avocado varieties was greater than 50%. Therefore, stress treatment at −4^°^C ∼ −6^°^C can be regarded as the boundary of relative electrical conductivity at 50%. Under low-temperature stress with relative electrical conductivity not higher than 50%, the magnitude of relative electrical conductivity can be used to roughly judge the level of cold resistance: the lower the relative electrical conductivity, the higher the cold resistance.

**Table 3:** Pearson correlation analysis between LT_50_ and relative electrical conductivity of 38 varieties of avocado leaves at different temperatures.

|  | Temperature |  |  |  |  |  |  |  |
| --- | --- | --- | --- | --- | --- | --- | --- | --- |
|  | CK | 4°C | 2°C | 0°C | -2°C | -4°C | -6°C | -8°C |
| Correlation coefficient | 0.269 | 0.420** | 0.649** | 0.727** | 0.708** | 0.415** | -0.151 | -0.124 |
| Sig | 0.102 | 0.009 | 0.000 | 0.000 | 0.000 | 0.010 | 0.365 | 0.593 |

\* Significant correlation at the 0.05 level (bilateral)
\*\* Significant correlation at the 0.01 level (bilateral)

### 2.4 Distribution law of the capacitance of 27 varieties of avocado leaves

As can be seen in Table 4, a large difference in capacitance of the same part was observed among different varieties of avocado leaves, and there was also a wide difference among different parts of the same leaf. The capacitance of the main vein at the leaf stalk base and middle leaf section was observed to be higher than that of the lateral vein at the leaf stalk base and middle leaf section, while the capacitance of the lateral vein at the leaf stalk base and middle leaf section was higher than that of the mesophyll. In about 92% of the 27 avocado varieties, the capacitance of the main vein at the leaf stalk base was significantly higher than that of the main vein middle leaf section; in about 96% of avocado varieties, there was no significant difference in capacitance between the lateral vein at the leaf stalk base and the lateral vein at the middle leaf section, nor was there a significant difference in the capacitance of the mesophyll in the leaf stalk base and middle leaf section for any avocado varieties; in about 63% of avocado varieties, there was no significant difference in capacitance among the main vein at the leaf tip, the lateral vein at the leaf tip and the mesophyll in the leaf tip. The capacitance of the main vein at the leaf stalk base, middle leaf section and leaf tip showed a significant decreasing trend, while there is no significant difference in capacitance between the lateral vein at the leaf stalk base and the lateral vein at the middle leaf section, but capacitance was significantly higher than the lateral vein at the leaf tip. In 58% of avocado varieties, there was no significant difference observed in the capacitance of mesophyll among the leaf stalk base, middle leaf section and leaf tip. This shows that there is a big difference in capacitance among different parts of avocado leaves, so when comparing the capacitance of different varieties, we should select the same part.

**Table 4:**
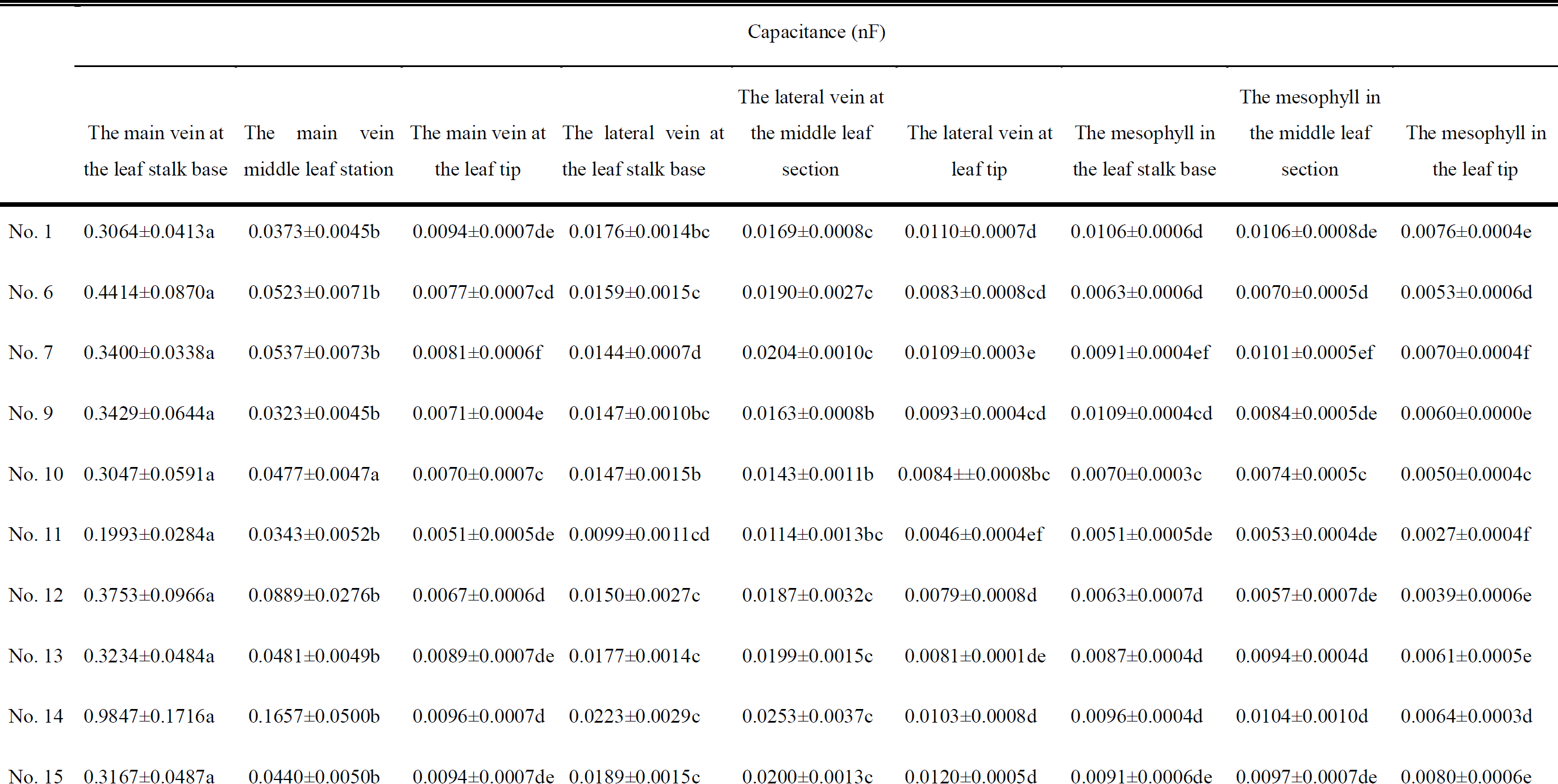
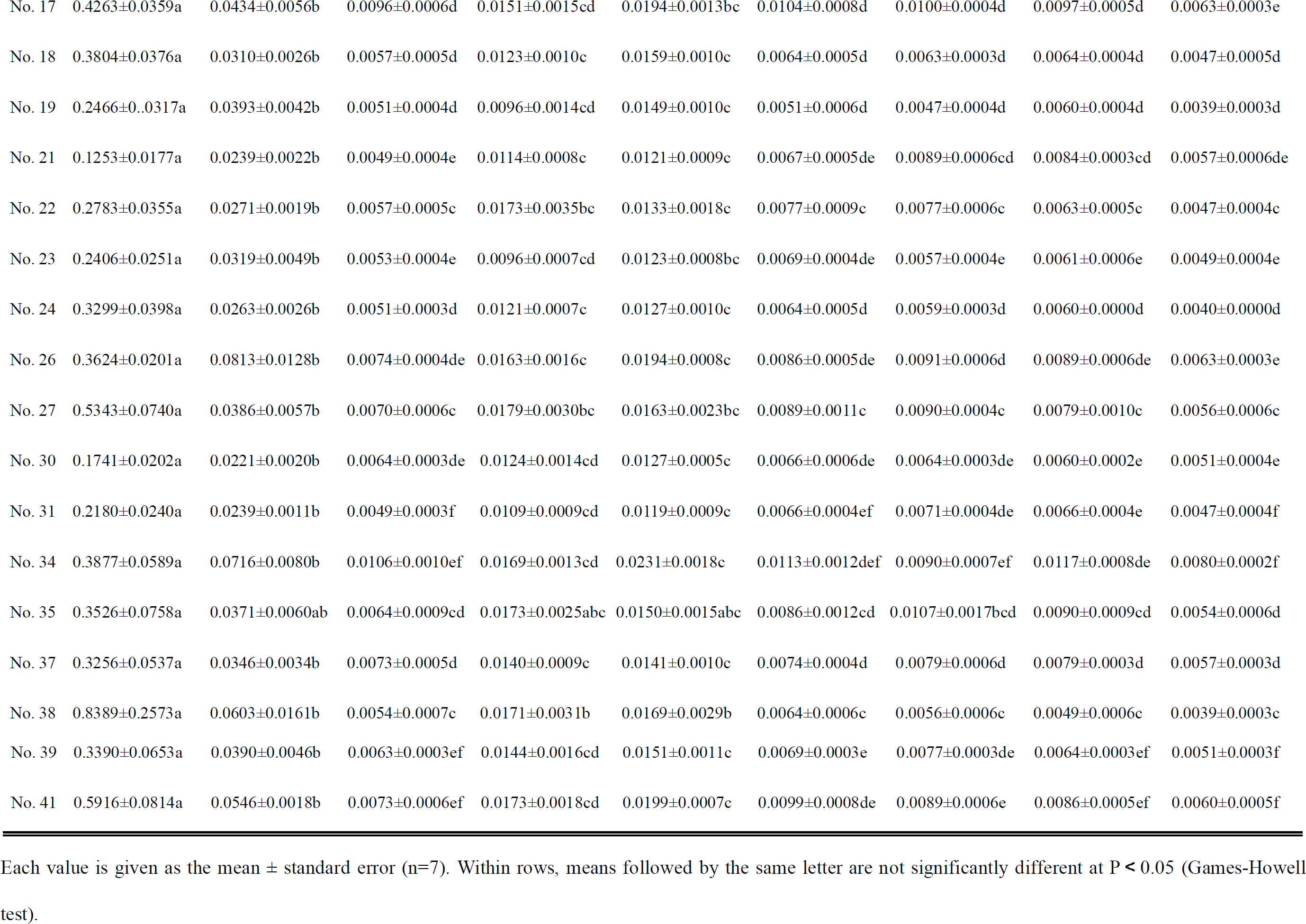
Capacitance of 26 varieties of avocado leaves at ambient temperature.

| Capacitance (nF) |  |  |  |  |  |  |  |  |  |
| --- | --- | --- | --- | --- | --- | --- | --- | --- | --- |
|  | The main vein at<br>the leaf stalk base | The main vein<br>middle leaf station | The main vein at<br>the leaf tip | The lateral vein at<br>the leaf stalk base | The lateral vein at<br>the middle leaf<br>section | The lateral vein at<br>leaf tip | The mesophyll in<br>the leaf stalk base | The mesophyll in<br>the middle leaf<br>section | The mesophyll in<br>the leaf tip |
| No. 1 | 0.3064±0.0413a | 0.0373±0.0045b | 0.0094±0.0007de | 0.0176±0.0014bc | 0.0169±0.0008c | 0.0110±0.0007d | 0.0106±0.0006d | 0.0106±0.0008de | 0.0076±0.0004e |
| No. 6 | 0.4414±0.0870a | 0.0523±0.0071b | 0.0077±0.0007cd | 0.0159±0.0015c | 0.0190±0.0027c | 0.0083±0.0008cd | 0.0063±0.0006d | 0.0070±0.0005d | 0.0053±0.0006d |
| No. 7 | 0.3400±0.0338a | 0.0537±0.0073b | 0.0081±0.0006f | 0.0144±0.0007d | 0.0204±0.0010c | 0.0109±0.0003e | 0.0091±0.0004ef | 0.0101±0.0005ef | 0.0070±0.0004f |
| No. 9 | 0.3429±0.0644a | 0.0323±0.0045b | 0.0071±0.0004e | 0.0147±0.0010bc | 0.0163±0.0008b | 0.0093±0.0004cd | 0.0109±0.0004cd | 0.0084±0.0005de | 0.0060±0.0000e |
| No. 10 | 0.3047±0.0591a | 0.0477±0.0047a | 0.0070±0.0007c | 0.0147±0.0015b | 0.0143±0.0011b | 0.0084±0.0008bc | 0.0070±0.0003c | 0.0074±0.0005c | 0.0050±0.0004c |
| No. 11 | 0.1993±0.0284a | 0.0343±0.0052b | 0.0051±0.0005de | 0.0099±0.0011cd | 0.0114±0.0013bc | 0.0046±0.0004ef | 0.0051±0.0005de | 0.0053±0.0004de | 0.0027±0.0004f |
| No. 12 | 0.3753±0.0966a | 0.0889±0.0276b | 0.0067±0.0006d | 0.0150±0.0027c | 0.0187±0.0032c | 0.0079±0.0008d | 0.0063±0.0007d | 0.0057±0.0007de | 0.0039±0.0006e |
| No. 13 | 0.3234±0.0484a | 0.0481±0.0049b | 0.0089±0.0007de | 0.0177±0.0014c | 0.0199±0.0015c | 0.0081±0.0001de | 0.0087±0.0004d | 0.0094±0.0004d | 0.0061±0.0005e |
| No. 14 | 0.9847±0.1716a | 0.1657±0.0500b | 0.0096±0.0007d | 0.0223±0.0029c | 0.0253±0.0037c | 0.0103±0.0008d | 0.0096±0.0004d | 0.0104±0.0010d | 0.0064±0.0003d |
| No. 15 | 0.3167±0.0487a | 0.0440±0.0050b | 0.0094±0.0007de | 0.0189±0.0015c | 0.0200±0.0013c | 0.0120±0.0005d | 0.0091±0.0006de | 0.0097±0.0007de | 0.0080±0.0006e |
| No. 17 | 0.4263±0.0359a | 0.0434±0.0056b | 0.0096±0.0006d | 0.0151±0.0015cd | 0.0194±0.0013bc | 0.0104±0.0008d | 0.0100±0.0004d | 0.0097±0.0005d | 0.0063±0.0003e |
| No. 18 | 0.3804±0.0376a | 0.0310±0.0026b | 0.0057±0.0005d | 0.0123±0.0010c | 0.0159±0.0010c | 0.0064±0.0005d | 0.0063±0.0003d | 0.0064±0.0004d | 0.0047±0.0005d |
| No. 19 | 0.2466±0.0317a | 0.0393±0.0042b | 0.0051±0.0004d | 0.0096±0.0014cd | 0.0149±0.0010c | 0.0051±0.0006d | 0.0047±0.0004d | 0.0060±0.0004d | 0.0039±0.0003d |
| No. 21 | 0.1253±0.0177a | 0.0239±0.0022b | 0.0049±0.0004e | 0.0114±0.0008c | 0.0121±0.0009c | 0.0067±0.0005de | 0.0089±0.0006cd | 0.0084±0.0003cd | 0.0057±0.0006de |
| No. 22 | 0.2783±0.0355a | 0.0271±0.0019b | 0.0057±0.0005c | 0.0173±0.0035bc | 0.0133±0.0018c | 0.0077±0.0009c | 0.0077±0.0006c | 0.0063±0.0005c | 0.0047±0.0004c |
| No. 23 | 0.2406±0.0251a | 0.0319±0.0049b | 0.0053±0.0004e | 0.0096±0.0007cd | 0.0123±0.0008bc | 0.0069±0.0004de | 0.0057±0.0004e | 0.0061±0.0006e | 0.0049±0.0004e |
| No. 24 | 0.3299±0.0398a | 0.0263±0.0026b | 0.0051±0.0003d | 0.0121±0.0007c | 0.0127±0.0010c | 0.0064±0.0005d | 0.0059±0.0003d | 0.0060±0.0000d | 0.0040±0.0000d |
| No. 26 | 0.3624±0.0201a | 0.0813±0.0128b | 0.0074±0.0004de | 0.0163±0.0016c | 0.0194±0.0008c | 0.0086±0.0005de | 0.0091±0.0006d | 0.0089±0.0006de | 0.0063±0.0003e |
| No. 27 | 0.5343±0.0740a | 0.0386±0.0057b | 0.0070±0.0006c | 0.0179±0.0030bc | 0.0163±0.0023bc | 0.0089±0.0011c | 0.0090±0.0004c | 0.0079±0.0010c | 0.0056±0.0006c |
| No. 30 | 0.1741±0.0202a | 0.0221±0.0020b | 0.0064±0.0003de | 0.0124±0.0014cd | 0.0127±0.0005c | 0.0066±0.0006de | 0.0064±0.0003de | 0.0060±0.0002e | 0.0051±0.0004e |
| No. 31 | 0.2180±0.0240a | 0.0239±0.0011b | 0.0049±0.0003f | 0.0109±0.0009cd | 0.0119±0.0009c | 0.0066±0.0004ef | 0.0071±0.0004de | 0.0066±0.0004e | 0.0047±0.0004f |
| No. 34 | 0.3877±0.0589a | 0.0716±0.0080b | 0.0106±0.0010ef | 0.0169±0.0013cd | 0.0231±0.0018c | 0.0113±0.0012def | 0.0090±0.0007ef | 0.0117±0.0008de | 0.0080±0.0002f |
| No. 35 | 0.3526±0.0758a | 0.0371±0.0060ab | 0.0064±0.0009cd | 0.0173±0.0025abc | 0.0150±0.0015abc | 0.0086±0.0012cd | 0.0107±0.0017bcd | 0.0090±0.0009cd | 0.0054±0.0006d |
| No. 37 | 0.3256±0.0537a | 0.0346±0.0034b | 0.0073±0.0005d | 0.0140±0.0009c | 0.0141±0.0010c | 0.0074±0.0004d | 0.0079±0.0006d | 0.0079±0.0003d | 0.0057±0.0003d |
| No. 38 | 0.8389±0.2573a | 0.0603±0.0161b | 0.0054±0.0007c | 0.0171±0.0031b | 0.0169±0.0029b | 0.0064±0.0006c | 0.0056±0.0006c | 0.0049±0.0006c | 0.0039±0.0003c |
| No. 39 | 0.3390±0.0653a | 0.0390±0.0046b | 0.0063±0.0003ef | 0.0144±0.0016cd | 0.0151±0.0011c | 0.0069±0.0003e | 0.0077±0.0003de | 0.0064±0.0003ef | 0.0051±0.0003f |
| No. 41 | 0.5916±0.0814a | 0.0546±0.0018b | 0.0073±0.0006ef | 0.0173±0.0018cd | 0.0199±0.0007c | 0.0099±0.0008de | 0.0089±0.0006e | 0.0086±0.0005ef | 0.0060±0.0005f |

Each value is given as the mean ± standard error (n=7). Within rows, means followed by the same letter are not significantly different at $P < 0.05$ (Games-Howell test).

### 2.5 Analysis of the correlation between the capacitance of different parts of avocado leaves and the LT_50_

Table 5 shows the results of a Pearson’s correlation analysis of the LT_50_ of 27 avocado varieties and the capacitance of the corresponding leaves. The LT_50_ of the various avocado varieties was significantly positively correlated with the capacitance of the main vein at the middle leaf section, the main vein at the leaf tip and the lateral vein at the leaf stalk base. Its correlation with the capacitance of the lateral vein at the leaf stalk base was at a very significant level. Although the correlation between the capacitance of other leaf parts and the LT_50_ was not significant at the level of 0.05, the correlation between the capacitance of the main vein at the leaf stalk base, lateral vein at the middle leaf section, mesophyll in the leaf stalk base and mesophyll in the middle leaf section and the LT_50_ was greater than 92%; the capacitance of the lateral vein at the leaf tip and the mesophyll in the leaf tip and the LT_50_ was around 80%. Thus we can conclude that the cold resistance of avocado varieties is significantly negatively correlated with the capacitance of the main vein at the middle leaf section, the main vein at the leaf tip and lateral vein at the middle leaf section of various varieties, and also negatively correlated with the capacitance of other parts of leaves. In other words, the higher the cold resistance, the lower the capacitance of leaf tissues.

**Table 5:** Pearson correlation analysis between the capacitance of different parts of avocado leaves and the LT50.

|  | Different parts of leaf |  |  |  |  |  |  |  |  |
| --- | --- | --- | --- | --- | --- | --- | --- | --- | --- |
|  | The main vein<br>at the leaf stalk<br>base | The main vein<br>middle leaf<br>station | The main vein<br>at the leaf tip | The lateral vein<br>at the leaf stalk<br>base | The lateral vein<br>at the middle<br>leaf section | The lateral<br>vein at leaf tip | The mesophyll<br>in the leaf stalk<br>base | The mesophyll<br>in the middle<br>leaf section | The mesophyll<br>in the leaf tip |
| Correlation<br>coefficient | 0.356 | 0.434* | 0.412* | 0.494** | 0.353 | 0.276 | 0.351 | 0.383 | 0.256 |
| Sig. | 0.074 | 0.027 | 0.036 | 0.010 | 0.077 | 0.172 | 0.079 | 0.054 | 0.206 |

\* Significant correlation at the 0.05 level (bilateral)
\*\* Significant correlation at the 0.01 level (bilateral)

### 2.6 Analysis of the capacitance of mango leaves with different levels of cold resistance

Table 6 shows the capacitance of the various parts of Dongmang and Tainong No. 1 leaves. The capacitance of nine parts of Dongmang leaves is lower than that of Tainong No. 1. According to the variance analysis in Table 7, the capacitance of the main vein in the middle leaf section of a Dongmang leaf is 0.189nF, significantly lower than that of Tainong No. 1, which is 0.3764nF; the capacitance of the main vein in a Dongmang leaf at stalk base and leaf tip is significantly lower than that of Tainong No. 1, while the cold resistance of Dongmang is higher than that of Tainong No. 1. This shows that the capacitance of some parts of mango leaves is negatively correlated with their cold resistance, and the value of capacitance can be used to identify the cold resistance of mango leaves. This is consistent with the results of capacitance research regarding the identification of avocado cold resistance.

**Table 6:** Capacitance of different parts of leaves in two mango plants.

|  | Capacitance (nF) |  |  |  |  |  |  |  |  |
| --- | --- | --- | --- | --- | --- | --- | --- | --- | --- |
|  | The main vein<br>at the leaf<br>stalk base | The main vein<br>middle leaf<br>station | The main vein<br>at the leaf tip | The lateral vein<br>at the leaf stalk<br>base | The lateral vein<br>at the middle<br>leaf section | The lateral<br>vein at leaf tip | The mesophyll<br>in the leaf<br>stalk base | The mesophyll<br>in the middle<br>leaf section | The mesophyll<br>in the leaf tip |
| Dongmang | 1.1394±0.1243 | 0.1890±0.0241 | 0.0065±0.0006 | 0.0037±0.0006 | 0.0043±0.0005 | 0.0033±0.0004 | 0.0023±0.0003 | 0.0020±0.0000 | 0.0020±0.0000 |
| Tainong No. 1 | 2.0448±0.3304 | 0.3764±0.0624 | 0.0117±0.0013 | 0.0039±0.0007 | 0.0047±0.0009 | 0.0037±0.0006 | 0.0025±0.0005 | 0.0023±0.0003 | 0.0023±0.0003 |

Each value is given as the mean ± standard error (n=10).

**Table 7:**
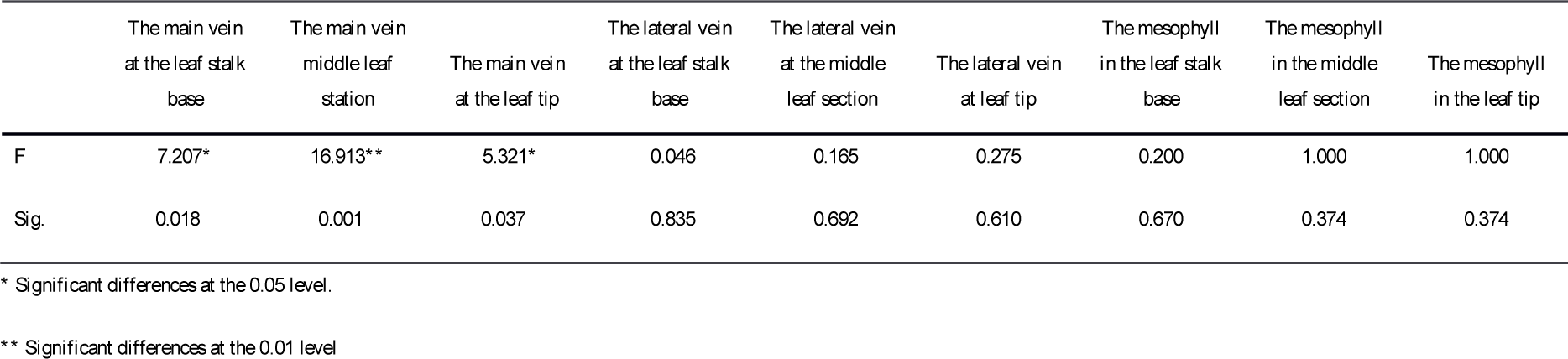
Variance analysis of counterpart of leaves in two mango plants.

## 3 Conclusions and Discussion

Temperature is an important ecological factor influencing the growth and development of plants (***Wei Jianxue, 2007***). Low temperature is one factor limiting the survival rate and distribution of plants, and when the temperature is lower than what is required for plant growth, plants will suffer with growth slowing or stopping (***Zhao Xijuan, 2013***), and some plants even dying (***Li Wenming, 2017***). Accurate, reliable identification and testing of plant cold resistance is essential for studies of the mechanism of chilling injury and cold resistance in plants, as well as breeding and innovation of excellent cold-resistant varieties (***Xu Chenxiang, 2012***). However, the methods used to research, evaluate and identify plant cold resistance are still limited by the species, organs, histologic types and physiological conditions of plants as well as research objectives and instruments (***Xu Chenxiang, 2014***).

Global scientific researchers have developed a few methods to research plant cold resistance, such as field production identification (***Sandra E. Vega et al., 1996; Zhang Qingfei et al., 2007; Zhao Xuemei et al., 2011***), freezing injury surveys (***Bao Wenjuan et al., 2005; Zhou Xihua et al., 2008***), manual frozen weather simulations (***Wei Jianxue, 2007; Liu Pin et al., 2009***), and mathematical modeling (***Timm is R et al., 1994; Zhang G et al., 2003***). However, existing plant cold resistance testing methods require manual freezing treatment, and most methods won’t produce results until after treatment at freezing temperature. So far no method is considered to be reliable and effective in testing plant cold resistance.

Previous research has shown that the permeability of a plant cell membrane changes under low-temperature stress (***Huang Yun et al., 2014***), and relative electrical conductivity—which can reflect the permeability of the plant cell membrane—can be measured to determine the level of injury from low temperatures, thereby providing a means to test plant cold resistance (***Zhu Genhai et al,, 1986***). On that basis, Rajashekar et al. used the logistic function curve to describe how low temperatures injured plant cell membranes, and proposed LT_50_ as the curve inflection, using it to decide plant cold resistance (***Rajashekar C et al., 1979***). The results of the present study show that the relative electrical conductivity of the 38 varieties of avocado leaves examined increases with lower stress temperatures, that there is a significant difference in relative electrical conductivity among different varieties at the same stress temperature (Table 1), and that there is also a big difference in LT_50_ among the different varieties (Table 2). This illustrates that relative electrical conductivity and LT_50_ can be used to effectively distinguish among the avocado varieties in terms of cold resistance. In addition, this study has also found that if the relative electrical conductivity is not greater than 50% under low-temperature stress, there is a significantly positive correlation between relative electrical conductivity and LT_50_, and if the relative electrical conductivity is greater than 50% under low-temperature stress, there is an insignificant correlation between relative electrical conductivity and LT_50_. Moreover, there is no absolute relationship between the electrical conductivity of a leaf at normal temperature and it’s sensitivity under low temperature(Table 1), so relative electrical conductivity at normal temperature is unsuitable for the identification of plant cold resistance

In this study, at normal temperature, a large difference was observed in the capacitance of the same part among different varieties of avocado leaves, and there was also a big difference in the capacitance of different parts of the same leaf (Table 4). The capacitance in the main vein at the middle leaf section, the main vein at the leaf tip and the lateral vein at the leaf stalk base was significantly positively correlated with LT_50_. The capacitance in the lateral vein at the leaf stalk base was found to have a particularly strong positive correlation with LT_50_ (Table 5). Thus we may conclude that the cold resistance of the avocado is significantly negatively correlated with the main vein at the middle leaf section, the main vein at the leaf tip and the lateral vein at the middle leaf section. In other words, the higher the cold resistance, the lower the capacitance of the various parts of leaves. This conclusion has been further verified in trials on two mango varieties with greatly different levels of cold resistance: the capacitance of the various parts of Dongmang leaves, which have high cold resistance, is lower than that of Tainong No. 1. In particular, the capacitance of the main vein at the leaf stalk base, middle leaf section, and leaf tip of Dongmang is significantly lower than in the corresponding regions of Tainong No. 1 (Tables 6 & 7). This might be because capacitance is a physical parameter that measures and represents the capacity of a capacitor (***D. Halliday et al., 2002***), and its magnitude relates to the state, structure and chemical composition of dielectrics. The tissue water in leaves is free water and a medium that contains charges. Within the range of unit volume, the higher the content of free water, the stronger the charge capacity, and the higher the capacitance (***Zhang Baishan, 2005; Wang Da, 2013***). However, the higher the content of free water, the lower cold resistance of leaf tissues (***Li Huimin and Lu Yan, 2013***).

Therefore, we argue that using a capacitance meter in the field it is practicable to identify a relationship in cold resistance among different varieties by determining the suitable parts of mature plant leaves. Moreover, this identification method does not demand plant samples undergo low-temperature stress, because it works at normal temperature, giving it more convenience, efficiency and non-destructiveness. However, this method can only be used to identify the relative cold resistance among different varieties, not to accurately determine the specific range of low temperatures to which a plant is resistant.

## Acknowledgments

The present study was carried out by our team under the guidance of Professor Li Shaopeng from the College of Tropical Agriculture and Forestry, Hainan University. We would like to thank all the team members for their hard work. The present study was accomplished with funding from the Hainan Provincial Ministry of Agriculture Bureau of Agricultural Reclamation as an experimental agricultural technology demonstration and service support project on avocado seedling breeding technology (151721301064071703-3)-and-avocado-resource-protection (151721301354051707-2).

## Appendix 1 Corresponding number of avocado varieties

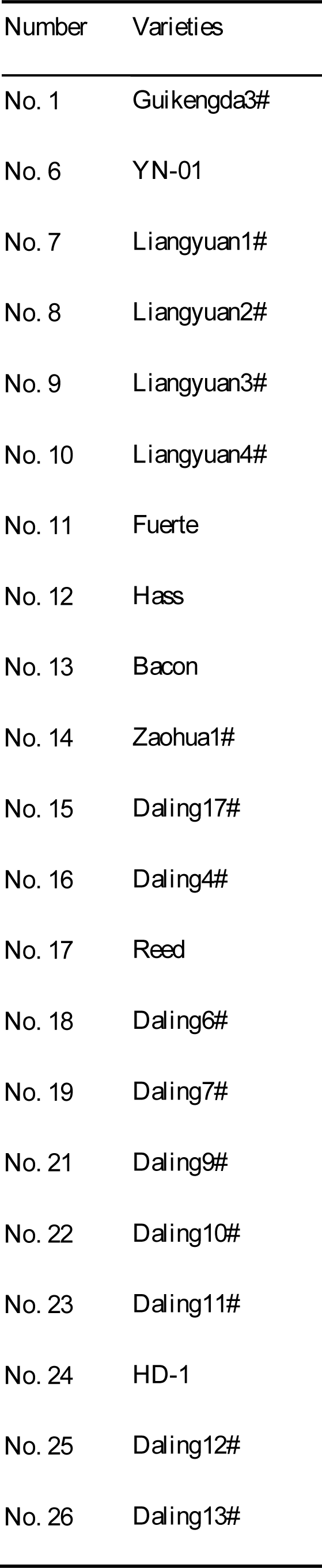
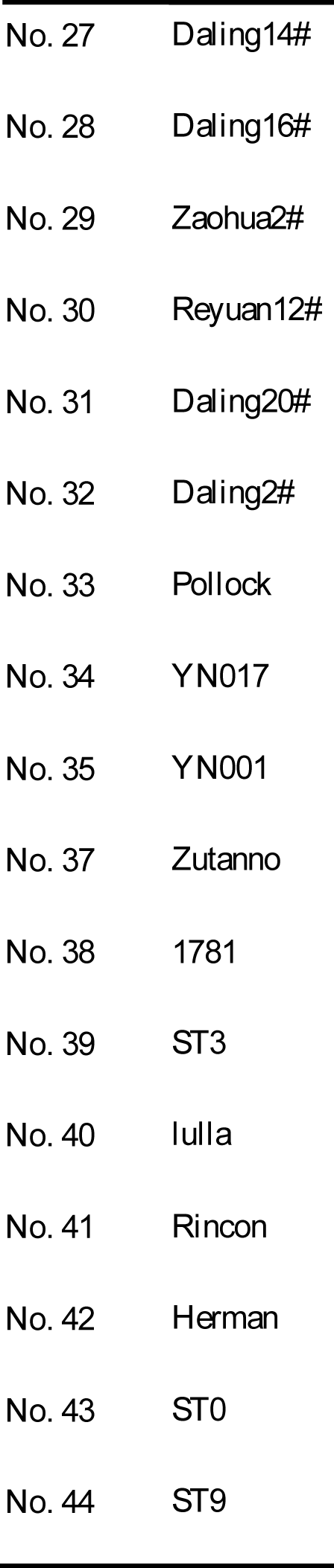

Appendix 1 Corresponding number of avocado varieties.
| Number | Varieties |
| --- | --- |
| No. 1 | Guikengda3# |
| No. 6 | YN-01 |
| No. 7 | Liangyuan1# |
| No. 8 | Liangyuan2# |
| No. 9 | Liangyuan3# |
| No. 10 | Liangyuan4# |
| No. 11 | Fuerte |
| No. 12 | Hass |
| No. 13 | Bacon |
| No. 14 | Zaohua1# |
| No. 15 | Daling17# |
| No. 16 | Daling4# |
| No. 17 | Reed |
| No. 18 | Daling6# |
| No. 19 | Daling7# |
| No. 21 | Daling9# |
| No. 22 | Daling10# |
| No. 23 | Daling11# |
| No. 24 | HD-1 |
| No. 25 | Daling12# |
| No. 26 | Daling13# |
| No. 27 | Daling14# |
| No. 28 | Daling16# |
| No. 29 | Zaohua2# |
| No. 30 | Reyuan12# |
| No. 31 | Daling20# |
| No. 32 | Daling2# |
| No. 33 | Pollock |
| No. 34 | YN017 |
| No. 35 | YN001 |
| No. 37 | Zutanno |
| No. 38 | 1781 |
| No. 39 | ST3 |
| No. 40 | lulla |
| No. 41 | Rincon |
| No. 42 | Herman |
| No. 43 | ST0 |
| No. 44 | ST9 |

